# D614G mutation of SARS-CoV-2 spike protein enhances viral infectivity

**DOI:** 10.1101/2020.06.20.161323

**Authors:** Jie Hu, Chang-Long He, Qing-Zhu Gao, Gui-Ji Zhang, Xiao-Xia Cao, Quan-Xin Long, Hai-Jun Deng, Lu-Yi Huang, Juan Chen, Kai Wang, Ni Tang, Ai-Long Huang

## Abstract

Coronavirus disease 2019 (COVID-19) is caused by severe acute respiratory syndrome coronavirus 2 (SARS-CoV-2). The spike (S) protein that mediates SARS-CoV-2 entry into host cells is a major target for vaccines and therapeutics. Thus, insights into its sequence variations are key to understanding the infection and antigenicity of SARS-CoV-2. A dominant mutational variant at position 614 of the S protein (aspartate to glycine, D614G mutation) was observed in the SARS-CoV-2 genome sequence obtained from the Nextstrain database. Using a pseudovirus-based assay, we identified that S-D614 and S-G614 protein pseudotyped viruses share a common receptor, human angiotensin-converting enzyme 2 (ACE2), which could be blocked by recombinant ACE2 with the fused Fc region of human IgG1. However, S-D614 and S-G614 protein demonstrated functional differences. First, S-G614 protein could be cleaved by serine protease elastase-2 more efficiently. Second, S-G614 pseudovirus infected 293T-ACE2 cells significantly more efficiently than did the S-D614 pseudovirus, especially in the presence of elastase-2. Third, an elastase inhibitor approved for clinical use blocked elastase-enhanced S-G614 pseudovirus infection. Moreover, 93% (65/70) convalescent sera from patients with COVID-19 could neutralize both S-D614 and S-G614 pseudoviruses with comparable efficiencies, but about 7% (5/70) convalescent sera showed reduced neutralizing activity against the S-G614 pseudovirus. These findings have important implications for SARS-CoV-2 transmission and immune interventions.

## Introduction

Severe acute respiratory syndrome coronavirus 2 (SARS-CoV-2) is a novel coronavirus reported in 2019 that caused the recent outbreak of coronavirus disease 2019 (COVID-19)^1^. By July 4, 2020, the World Health Organization (WHO) reported that 10.92 million people worldwide had been infected with SARS-CoV-2, and that 523 000 individuals had died of COVID-19. This pandemic has had a significant adverse impact on international, social, and economic activities. Coronaviruses are enveloped, positive-stranded RNA viruses that contain the largest known RNA genomes to date. The RNA genome of SARS-CoV-2 has been rapidly sequenced to facilitate diagnostic testing, molecular epidemiologic source tracking, and development of vaccines and therapeutic strategies^2^. The mutation rate for RNA viruses is extremely high, which may contribute to their transmission and virulence. The only significant variation in the SARS-CoV-2 spike (S) protein is a non-synonymous D614G (aspartate (D) to glycine (G)) mutation^3^. Primary data showed that S-G614 is a more pathogenic strain of SARS-CoV-2 with high transmission efficiency^3^. However, whether the D614G mutation in the S protein affects viral entry and infectivity in a cellular model is still unclear.

The S protein of coronavirus, the major determinant of host and tissue tropism, is a major target for vaccines, neutralizing antibodies, and viral entry inhibitors^4,5^. Similar to SARS-CoV, the cellular receptor of SARS-CoV-2 is angiotensin-converting enzyme 2 (ACE2); however, the SARS-CoV-2 S protein has a 10- to 20-fold higher affinity for ACE2 than the corresponding S protein of SARS-CoV^6,7^. Coronaviruses use two distinct pathways for cell entry: protease-mediated cell-surface and endosomal pathways^8^. The S proteins of several coronaviruses are cleaved by host proteases into S1 subunit for receptor binding and S2 subunit for membrane fusion at the entry step of infection. Several cellular proteases including furin, transmembrane protease serine 2 (TMPRSS2), and cathepsin (Cat) B/L are critical for priming the SARS-CoV-2 S protein to enhance ACE2-mediated viral entry^4^. Recently, Bhattacharyya et al. (2020) reported that a novel serine protease (elastase-2) cleavage site was introduced into the S-G614 protein of SARS-CoV-2^9^. However, it is unknown whether the S-G614 protein can be processed and activated by elastase-2 in a cellular model. The S protein plays a key role in the evolution of coronaviruses to evade the host immune system. It is still uncertain whether the D614G mutation affects the antigenic properties of the S protein. Meanwhile, whether elastase-2 inhibitors and convalescent serum samples of COVID-19 can block the infection of the D614G variant of SARS-CoV-2 remains unknown.

In this study, we analyzed the S gene sequences of SARS-CoV-2 submitted to the Global Initiative on Sharing All Influenza Data (GISAID) database. We examined the expression and cleavage of S-D614 and S-G614 protein in cell lines. Using a luciferase (Luc)-expressing lentiviral pseudotype system, we established a quantitative pseudovirus-based assay for the evaluation of SARS-CoV-2 cell entry mediated by the viral S protein variants. We also compared the neutralizing sensitivity of the S-D614 and S-G614 protein pseudoviruses to convalescent sera from patients with COVID-19. Our study provides further insights into the transmission and immune interventions of this newly emerged virus.

## Results

### D614G mutation of SARS-CoV-2 S protein was globally distributed

The S protein of SARS-CoV-2, which contains 1,273 amino acids, forms a trimeric spike on the virion surface and plays an essential role in viral entry. We analyzed the SARS-CoV-2 S protein amino acid sequence from viral genomic sequences in the GISAID database. In line with prior reports, we found a globally distributed S protein mutation, D614 to G614, which represented 64.6% of all the analyzed sequences (Fig. 1 and Table 1). Among the top 10 most abundant non-synonymous mutations observed in the S protein, the relative abundance of the D614G mutant (the clade G) was the highest around the world, indicating that G614 strain may be selectively advantageous. Since the S protein is critical to coronavirus infection, we sought to explore the potential impact of the most prevalent D614G mutation on the S protein structure, expression, and function. Using the cryo-electron microscopy structure of S protein (PBD ID: 6ZGE) determined by Wrobel et al.^10^, we analyzed the potential effects of the D614G mutation. As shown in Fig. 1b, residue D614 is located at the C-terminal region of the S1 domain, which directly associates with S2. On the one hand, residue D614 forms a hydrogen bond with backbone of G593 and a salt bridge with K854, and then is capped by the folded 833-855 motif of chain B (Fig. 1c). D614G mutation would abolish these interactions. This change may destabilize the locked conformation to promote receptor binding domain (RBD) opening and potentially increase virus-receptor binding and membrane fusion activities. On the other hand, we observed that D614 is remarkably close to the N-linked glycosylation site N616 (Fig 1c). Thus, the D614G mutation may enhance the fitness of SARS-CoV-2 by increasing S protein stability and participating in glycosylation. Further studies are required to determine the structure of S-G614 protein.

**Fig. 1.**
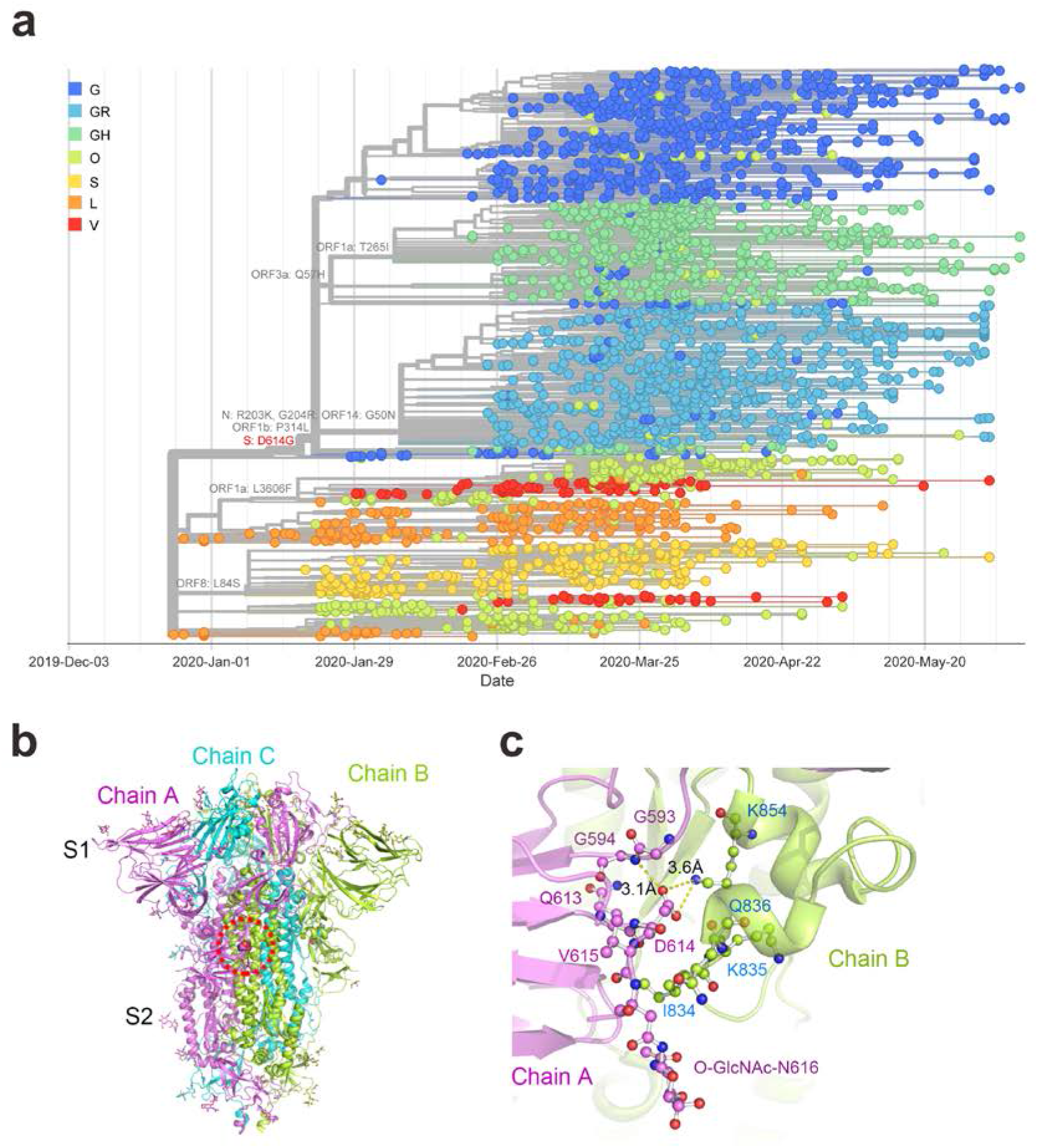
Phylogenetic timetree of SARS-CoV-2 and structural features of its spike (S) protein. **a** Phylogenetic analysis of SARS-CoV-2 sequences from GISAID database. Prevalence of S-D614G genomes over time produced by the Nextstrain analysis tool using GISAID dataset (*n* = 2,834 genomes samples from January 2020 to May 2020). **b** Cartoon representation of the trimeric SARS-CoV-2 spike structure (PBD ID: 6ZGE). Three spike chains are colored in violet, lemon, and aquamarine, respectively. D614 is located at the C-terminal of the S1 subunit and is shown as violet spheres in the red dashed circle. **c** D614 involves the interactions between two S chains. Residues are highlighted by violet or lemon spheres and sticks. Yellow dashed lines indicate hydrogen bonds or salt bridges. PyMOL software (Schrödinger, LLC) was used to generate all rendered structural images.

**Table 1.** Mutations in spike protein of SARS-CoV-2.

| **Mutations in Spike Protein** | **No. of mutation** | **No. of wildtype** | **Total No. of sequences** | **Mutation (%)** |
| --- | --- | --- | --- | --- |
| **D614G** | 2995 | 1637 | 4632 | 64.659 |
| **A829T** | 37 | 4602 | 4639 | 0.798 |
| **L5F** | 33 | 4614 | 4647 | 0.710 |
| **H146Y** | 26 | 4594 | 4620 | 0.563 |
| **P1263L** | 15 | 4631 | 4646 | 0.323 |
| **V483A** | 12 | 4387 | 4399 | 0.273 |
| **S939F** | 12 | 4626 | 4638 | 0.259 |
| **R78M** | 10 | 4623 | 4633 | 0.216 |
| **E583D** | 9 | 4639 | 4648 | 0.194 |
| **A845S** | 9 | 4632 | 4641 | 0.194 |

Top 10 abundant non-synonymous mutations observed in S protein of SARS-CoV-

### D614G mutation enhanced the cleavage of S protein variant by proteases

As the D614G mutation is proximal to the S1 cleavage domain, we predicted potential cleavage sites of proteases in S protein variants using PROSPER^11^, and identified a novel serine protease (elastase-2) cleavage site at residues 615-616 on the S1-S2 junction of the S-G614 protein (Fig. 2a, Supplementary information, Table S2). To evaluate the expression and cleavage of SARS-CoV-2 S protein in a human cell line, the codon-optimized S protein-expressing plasmids (pS-D614 and pS-G614) were transfected into HEK 293T cells. The immunoblot analysis of whole cell lysates revealed that both S-D614 and S-G614 proteins showed two major protein bands (unprocessed S and cleaved S1 subunit), when allowed to react with the monoclonal antibody targeting the RBD on the SARS-CoV-2 S protein (Fig. 2b). However, the pS-G614-transfected cells showed a stronger S1 signal than pS-D614-transfected cells, indicating that the D614G mutation altered the cleavability of the S protein by cellular proteases. Moreover, the elastase inhibitor sivelestat sodium significantly decreased the S1 signal of S-D614 protein (Fig. 2b). These data indicate that the D614G mutation of SARS-CoV-2 S facilitates its cleavage by host serine protease elastase-2. The coronavirus S protein must be cleaved by host proteases to enable membrane fusion, which is critical for viral entry. Next, we sought to explore the impact of D614G mutation on viral entry.

**Fig. 2.**
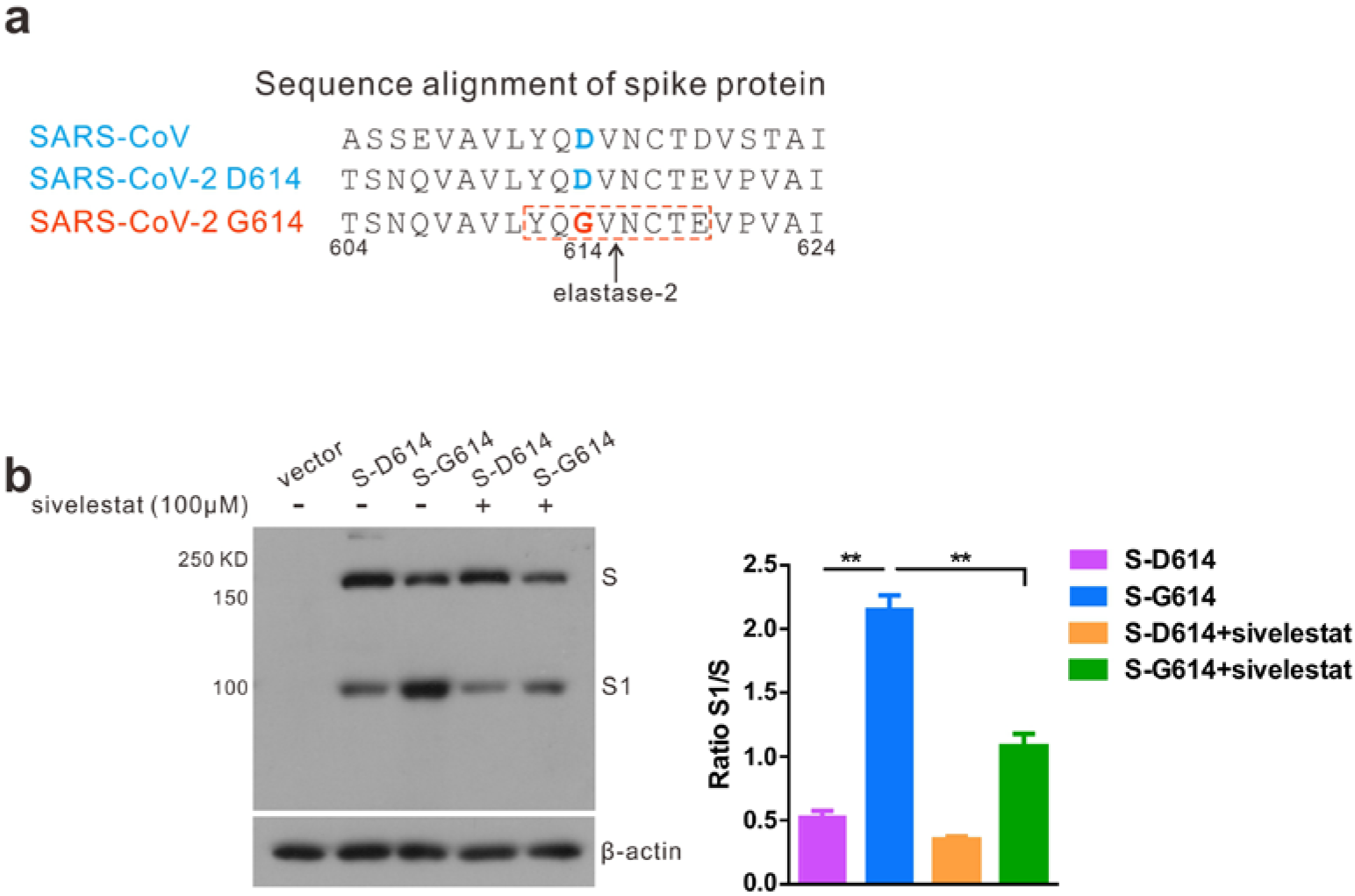
Detection of S-D614 and S-G614 protein expression and cleavage. **a** An additional serine protease (elastase-2) cleavage site in S1-S2 junction was identified in S-614G protein of SARS-CoV-2. **b** Detection of S protein expression in HEK 293T cells by Western blot using the anti-RBD (receptor-binding domain) monoclonal antibody. Cells were transfected with pS-D614 or pS-G614 plasmids or with an empty vector and incubated with or without sivelestat sodium. To compare the S1 and S ratio, integrated density of S1/S was quantitatively analyzed using ImageJ software. n = 3, ±SD. \*\**P* < 0.01.

### Evaluating viral entry efficacies between S-D614 and S-G614 pseudotyped lentiviral particles

Lentiviral vectors can be pseudotyped with various heterologous viral glycoproteins that modulate cellular tropism and entry properties^12^. Due to the highly pathogenic nature of SARS-CoV-2, infectious SARS-CoV-2 must be handled in a biosafety level 3 (BSL-3) facility. We generated pseudotyped SARS-CoV-2 based on the viral S protein using a lentiviral system, which introduced a Luc (luciferase) reporter gene for quantification of SARS-CoV-2 S-mediated entry. Thereafter, pNL4-3.Luc.R-E-was co-transfected with pS-D614 and pS-G614 to package the SARS-CoV-2 S pseudotyped single-round Luc virus in HEK 293T cells.

The titers of S-D614 and S-G614 protein pseudotyped viruses were determined by reverse transcriptase quantitative polymerase chain reaction (RT-qPCR) expressed as the number of viral RNA genomes per mL, and then adjusted to the same concentration (3.8 × 10^4^ copies in 50 μL) for the following experiments. The virus infectivity was determined by a Luc assay expressed as relative luminescence units (RLU). HEK 293T cells expressing human ACE2 (293T-ACE2) were used to test the correlation between ACE2 expression and pseudoviral susceptibility. The vesicular stomatitis virus G (VSV-G) pseudovirus was used as control. As shown in Fig. 3a, both HEK 293T and 293T-ACE2 cells could be effectively transduced by VSV-G pseudovirus. However, the entry of S-D614 and S-G614 pseudoviruses is highly dependent on its cellular receptor ACE2 expression. The 293T-ACE2 cells showed an approximately 250-fold and 530-fold increase in Luc activity when transduced by S-D614 and S-G614 pseudoviruses compared to HEK 293T cells, respectively (Fig. 3a). We then detected the inhibitory ability of ACE2-Ig, a fusion protein consisting of the extracellular domain (Met 1-Ser 740) of human ACE2 linked to the Fc region of human IgG1 at the C-terminus^13^. Both S-D614 and S-G614 pseudoviruses were potently inhibited by ACE2-Ig, and the IC_50_ (the concentration causing 50% inhibition of pseudoviral infection) values were 0.13 and 0.15 μg/mL, respectively (Fig. 3b). To further compare the viral entry efficiency meditated by S variants, we detected the Luc activity at different time points post-infection. With the G614 S variant, the increase in viral transduction over the D614 variant was 2.2-fold at 48 h post-infection. The highest transduction efficiency (approximately 2.5 × 10^4^ RLU) was observed 72 h post-infection with the S-G614 pseudovirus, which was approximately 2.4-fold higher than that of the S-D614 pseudovirus (Fig. 3c). These data suggest that the D614G mutation in S protein significantly promotes viral entry into ACE2-expressing cells, and ACE2-Ig efficiently blocks both wild-type and mutant S pseudotype virus infection.

**Fig. 3.**
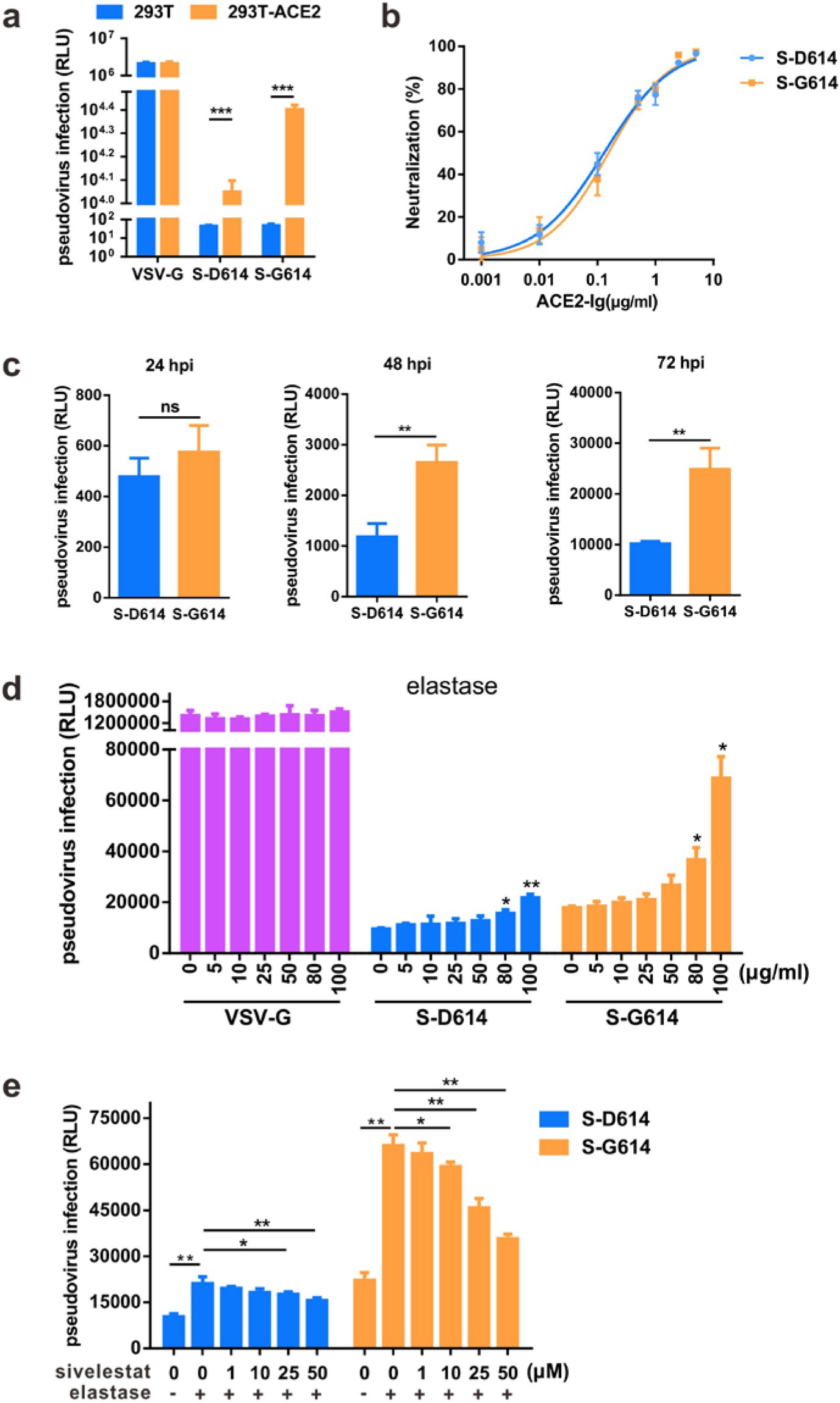
The S-G614 protein pseudotyped virus showed increased infectivity. **a** HEK 293T and 293T-ACE2 (human angiotensin-converting enzyme 2) cells were infected with lentiviruses pseudotyped with VSV-G and SARS-CoV-2 S protein variants. Virus titers were quantified by RT-qPCR and adjusted to 3.8 × 10^4^ copies in 50 μL to normalize input virus doses. The relative luminescence units (RLU) detected 72 h post-infection (hpi). **b** Inhibition of pseudoviral entry by ACE2-Ig. Pseudoviruses were pre-incubated with ACE2-Ig and added to 293T-ACE2 cells, then RLU was measured at 72 hpi. **c** Viral entry efficiency meditated by S variants. The RLU was measured at 24-72 hpi. **d-e** D614G mutation facilitates elastase-2 induced pseudoviral entry. 293T-ACE2 cells were treated with elastase for 5 min and then infected with pseudotyped viruses containing the S-D614 or S-G614 mutant in the presence of various concentrations of sivelestat sodium. RLU was measured at 72 hpi. n = 3, ±SD. \**P* < 0.05, \*\**P* < 0.01. ns, not significant.

We next explored the mechanism by which S-G614 increased pseudoviral infectivity. The proteolytic activation of S protein is required for coronavirus infectivity, and the protease-mediated cell-surface pathway is critical for SARS-CoV-2 entry^4^. Since we observed S-G614 could be more efficiently cleaved by the host protease when exogenously expressed in 293T cells, we assumed that host proteases may be involved in the enhancement of S-G614 viral entry. As shown in Fig. 3d, when 293T-ACE2 cells were treated with 100 μg/mL elastase before virus infection, the RLU value in S-G614 pseudovirus-infected cells (6.9 × 10^4^) was about 3.1 fold higher than that of the S-D614 (2.2 × 10^4^), indicating that residue G614 facilitates elastase-induced viral entry. In addition, the clinically proven serine protease inhibitor sivelestat sodium, which is active against elastase, dose-dependently blocked S-G614-driven entry into 293T-ACE2 in the presence of 100 μg/mL elastase (Fig. 3e). These data indicated that the infectivity of S-G614 pseudovirus containing an additional elastase-2 cleavage site is enhanced by exogenous elastase; therefore, S-G614 pseudovirus is more sensitive to sivelestat sodium than the S-D614 pseudovirus.

We also tested two other protease inhibitors, camostat mesylate and E-64d, which block host TMPRSS2 and CatB/L, respectively. In 293T-ACE2 cells lacking TMPRSS2, the serine protease inhibitor camostat mesylate did not inhibit S-D614 or S-G614 pseudoviral infection (Fig. S1a), while the cysteine protease inhibitor E-64d significantly blocked the entry of these two pseudoviruses, with IC_50_ values of 0.37 μM and 0.24 μM for S-D614 and S-G614 pseudoviruses, respectively (Fig. S1b). As expected, these protease inhibitors had no impact on VSV-G pseudovirus infection. Together, these results suggest that S-meditated viral entry into 293T-ACE2 cells deficient in TMPRSS2 is endosomal cysteine protease CatB/L-dependent; therefore, S-D614 and S-G614 pseudoviruses showed similar sensitivity to the CatB/L inhibitor E-64d in 293T-ACE2 cells.

### Neutralization effect of convalescent sera from patients with COVID-19 against S-D614 and S-G614 pseudoviruses

Neutralizing antibodies are important for prevention of and possible recovery from viral infections. However, as viruses mutate during replication and spread, host neutralizing antibodies generated in the earlier phase of the infection may not be as effective later on^14,15^. To test whether D614G mutations could affect the neutralization sensitivity of the virus, the neutralization activity of serum samples from convalescent patients with COVID-19 against SARS-CoV-2 S-D614 and S-G614 pseudoviruses were evaluated. To perform the neutralization assay, 50 μL pseudovirus (3.8 × 10^4^ copies) was incubated with serially diluted sera. As shown in Fig. 4a, the inhibition rate of sera from convalescent COVID-19 patients was analyzed at a single dilution of 1:1000, among the 70 tested sera, 65 of them showed neutralizing activities against both S-D614 and S-G614 pseudoviruses with comparable efficiencies. However, five sera samples (patients 1#, 7#, 40#, 42# and 52#) showed decreased inhibition rate against the S-G614 pseudovirus. The serum from patient 1# failed to neutralize the S-G614 pseudovirus, even though it neutralized about 30% of the S-D614 pseudovirus at a 1:1000 dilution. Then, the inhibition curves and half-maximal inhibitory dose (ID_50_) of sera samples from five convalescent patients and one healthy donor were analyzed. Sera from patients 17# and 39#, which were able to neutralize both pseudoviruses to similar degrees, showed similar ID_50_ values (Fig. 4b). However, sera from patients 1#, 7#, 40#, 42# and 52# showed relative high neutralizing activity against the S-D614 pseudovirus with an ID_50_ ranging from 729 to 1524, but showed decreased neutralizing activity against the S-G614 pseudovirus with an ID_50_ ranging from 216 to 367, indicating a 2.5- to 5.9-fold reduction in neutralizing titers (Fig. 4b). These data indicate that most antisera (93%) from patients, likely infected with earlier SARS-CoV-2 variant, can cross neutralize S-G614 variant, although D614G mutation decreases the neutralization sensitivity to 7% convalescent sera.

**Fig. 4.**
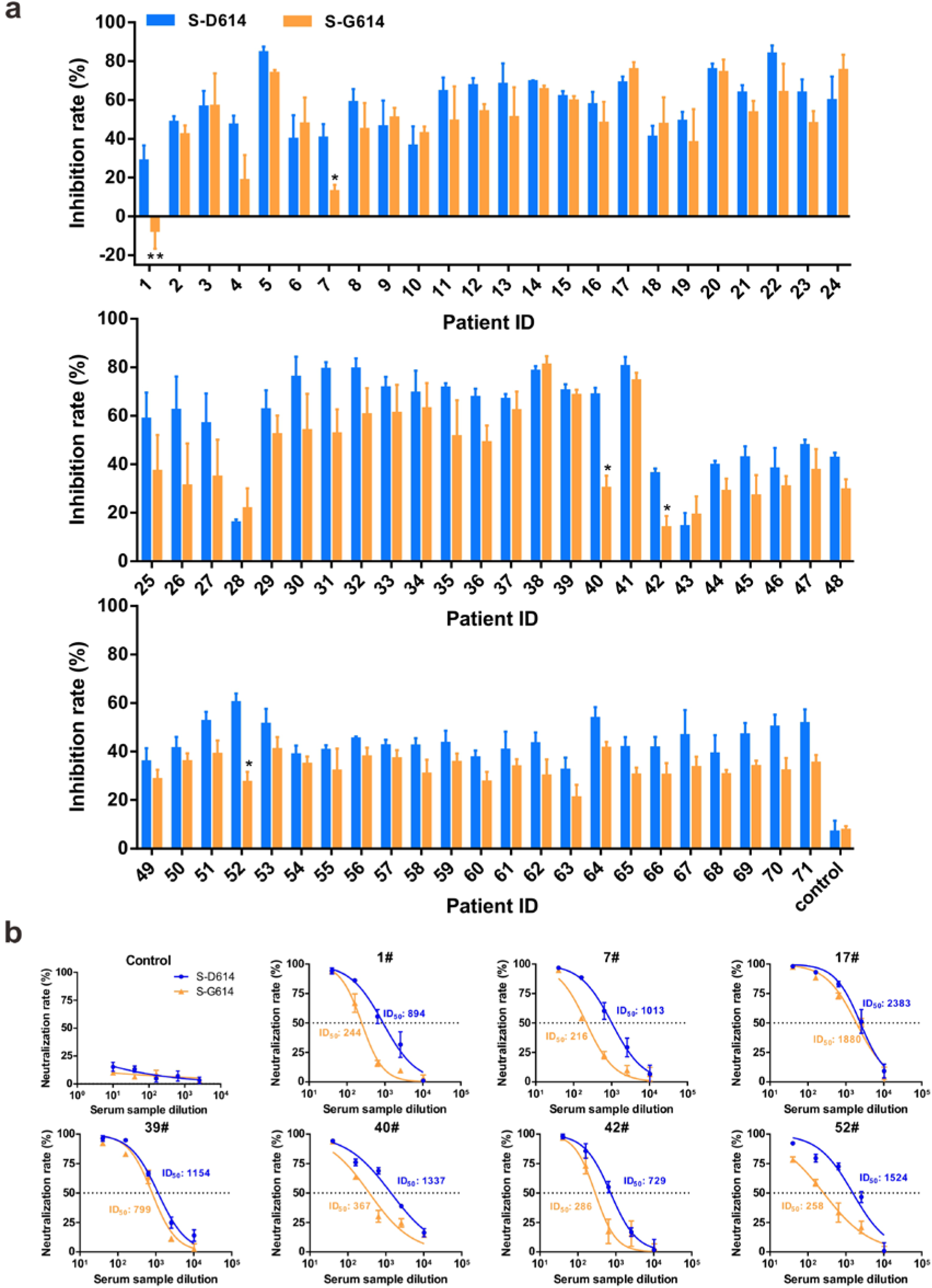
Detection of neutralizing antibodies in convalescent sera against spike (S)-D614 and S-G614 protein pseudotyped viruses. **a** The inhibition rate of convalescent sera from 70 patients with COVID-19 against S-D614 and S-G614 pseudoviruses. A serum sample from healthy individual was tested as a negative control. Convalescent sera were collected 2–4 weeks after symptom onset from confirmed case patients (1#–70#). Sera was analyzed at a single dilution of 1:1000. RLU was measured at 72 hpi. We then compared the RLU of serum neutralized sample to the control and calculated the inhibition rate. n = 3, ±SD. **b** The inhibition curves for serum samples from seven convalescent patients and a healthy donor. The initial dilution was 1:40, followed by 4-fold serial dilution. Neutralization titers were calculated as 50% inhibitory dose (ID_50_), expressed as the serum dilution at which RLUs were reduced by 50% compared with virus control wells after subtraction of background RLU in cell control wells.

## Discussion

Pseudovirus-based assays have been widely used for the study of cellular tropism, receptor recognition, viral inhibitors, and evaluation of neutralizing antibodies without the requirement of BSL-3 laboratories. We constructed pseudotyped SARS-CoV-2 based on the viral S protein using a lentiviral system, which incorporated a Luc reporter gene for easy quantification of coronavirus S protein-mediated entry. We investigated the major mutation in the S protein at position 614 and found that serine protease elastase-2 participates in the proteolytic activation of the S-G614 protein, thereby enhancing viral entry into 293T-ACE2 cells (Fig. 5). We demonstrated the potential role of the elastase-2 inhibitor sivelestat in blocking S-G614 pseudovirus infection in the presence of elastase. Although S-D614 and S-G614 variants could be similarly neutralized by most antisera, we found that the S-G614 pseudovirus was more resistant to some neutralizing antisera than S-D614 pseudovirus.

**Fig. 5.**
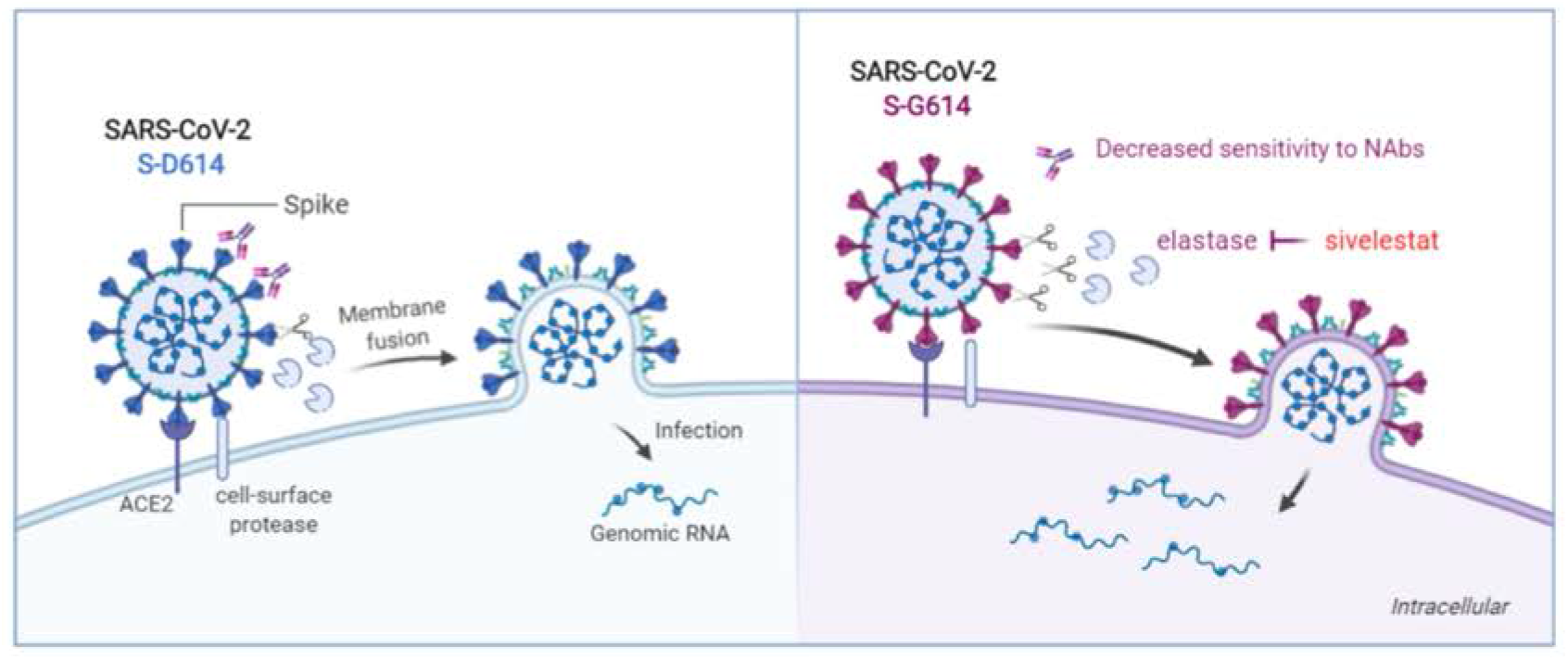
Proposed working model of this study. Protease-mediated cell-surface pathway is critical for SARS-CoV-2 entry and infection. D614G mutation introduces an additional elastase-2 cut site in the spike protein of SARS-CoV-2, thereby promoting its cleavage and enhancing viral entry into host cells. The elastase-2 inhibitor sivelestat partially blocks S-G614 pseudovirus infection in the presence of elastase. Both S-D614 and S-G614 variants could be similarly neutralized by most antisera, however, S-G614 mutant is more resistant to neutralizing antibodies (NAbs) from some patients infected with earlier SARS-CoV-2 variant.

Several findings stand out in our study: First, we found that the entry efficiency of S-G614 pseudotyped virus was about 2.4 times higher than that of the S-D614 pseudovirus when viral input doses were normalized, suggesting that D614G mutation promotes the infectivity of SARS-CoV-2 and enhances viral transmissibility. Since the pseudoviruses were employed in a single round of infection assay, this seemingly small increase in entry activity could cause a large difference in viral infectivity *in vivo*. Yao et al. (2020) reported that a patient-derived viral isolate ZJU-1, which harbors the D614G mutation, has a viral load 19 times higher than isolate ZJU-8 (harboring the S-D614) when Vero-E6 cells were infected^16^. However, the ZJU-1 isolate contained two other non-synonymous mutations in open reading frame 1a (ORF1a) and envelope (E) gene. Our results also elucidated the cause of increased entry efficiency due to the D614G mutation in the S protein of SARS-CoV-2. The S-G614 protein contains a novel serine protease cleavage site, so it could be cleaved by host elastase-2 more efficiently. Previously studies on SARS-CoV demonstrated that the protease-mediated cell surface entry facilitated a 100- to 1,000-fold increase in efficient infection compared to the endosomal pathway in the absence of proteases^17^.

Second, we found that elastase-enhanced S-G614 pseudoviral infection could be partially blocked by sivelestat sodium. Elastase-2, also known as neutrophil elastase, plays an important role in degenerative and inflammatory diseases. Sivelestat, a drug approved to treat acute respiratory distress syndrome (ARDS) in Japan and South Korea, has a beneficial effect on the pulmonary function of patients with ARDS and systemic inflammatory response syndrome^18^. About 10– 15% of patients with COVID-19 progress to ARDS^19^. Since sivelestat may not only mitigate the damage of neutrophil elastase on lung connective tissue, but also limit virus spread by inhibiting S protein processing, Mohamed et al. (2020) advocated the use of sivelestat to alleviate neutrophil-induced damage in critically ill patients with COVID-19^20^. Our *in vitro* results also indicate that sivelestat sodium or similar neutrophil elastase inhibitors might be an effective option for treatment of COVID-19 caused by SARS-CoV-2 harboring the D614G mutation.

Third, we observed that the S-G614 pesudovirus was more resistant to neutralization by convalescent sera from patients, likely infected in mid- to late-January when wild-type (D614) virus was mainly circulating in China. Koyama et al. (2020) reported that D614G is located in one of the predicted B-cell epitopes of SARS-CoV-2 S protein, and this is a highly immunodominant region and may affect the effectiveness of vaccine with wild-type S protein^14^. D614 is conserved in the S protein of SARS-CoV in 2003. Previous studies of SARS-CoV suggested that the peptide S597–625 is a major immunodominant peptide in humans and elicits a long-term B-cell memory response after natural infection with SARS-CoV^21^. Regions between amino acids 614 and 621 of SARS-CoV-2 S protein were also identified as a B-cell epitope by different methods, and the change in D614G may affect the antigenicity of this region^22^. In our study, we observed that 7% (5/70) convalescent sera showed markedly different neutralization activities between S-G614 and S-G61 protein pseudotyped viruses, indicating that the D614G mutation reduces the sensitivity to neutralizing antibodies from some patients infected with earlier SARS-CoV-2 variant. Whether these patients were at high risk of reinfection with the S-G614 variant should be explored in further studies. It will also be important to determine the breadth of the neutralizing capacity of vaccine-induced neutralizing antibodies.

Recently, several groups also reported that S-G614 enhances viral infectivity based on pseudovirus assays^23–26^, but due to the small sample size, they found no effect on the neutralization sensitivity of the virus^23,24,26^. Given the evolving nature of the SARS-CoV-2 RNA genome, antibody treatment and vaccine design require further considerations to accommodate D614G and other mutations that may affect the immunogenicity of the virus.

Our study had some limitations. First, 19 amino acids of S protein on the C-terminal were not included in order to improve the packaging efficiency of the SARS-CoV-2 S protein pseudotyped virus, and this pseudovirus only recapitulates viral entry events. Therefore, additional assays with authentic SARS-CoV-2 viruses are required. Second, we only tested neutralizing antibodies against the S protein. Previous studies on SARS-CoV indicated that only a small fraction of memory B cells specific for SARS-CoV antigens are directed against neutralizing epitopes present on the S protein^27^. Third, in addition to D614G, further studies on other mutations in the S protein are needed to evaluate their impact on SARS-CoV-2 infectivity, pathogenicity, and immunogenicity. Further studies are needed to determine the impact of these mutations on the severity of COVID-19.

In summary, we established a SARS-CoV-2 S protein-mediated pseudoviral entry assay and explored the cellular entry of S-D614 and S-G614 pseudotyped viruses. Our study provided evidence that the D614G mutation introduces an additional elastase-2 cut site in the S protein, thereby promoting its cleavage and viral cell entry, resulting in SARS-CoV-2 becoming more infectious. Importantly, the D614G mutation reduced the sensitivity of the virus to serum neutralizing antibodies in 7% convalescent patients with COVID-19. Our study will be helpful for understanding SARS-CoV-2 transmission and for the design of vaccines and therapeutic interventions against COVID-19.

## Materials and Methods

### Plasmids

The codon-optimized gene encoding SARS-CoV-2 S protein (GenBank: QHD43416) with C-terminal 19-amino acid deletion was synthesized by Sino Biological Inc (Beijing, China), and cloned into the *Kpn*I and *Xba*I restriction sites of pCMV3 vector (pCMV3-SARS-CoV-2-S-C19del, denoted as pS-D614). The D614G mutant S-expressing plasmid (denoted as pS-G614) was constructed by site-directed mutagenesis, with pS-D614 plasmid as a template. The HIV-1 NL4-3 ΔEnv Vpr luciferase reporter vector (pNL4-3.Luc.R-E-) constructed by N. Landau^28^ was provided by Prof. Cheguo Cai from Wuhan University (Wuhan, China). The VSV-G-expressing plasmid pMD2.G was provided by Prof. Ding Xue from Tsinghua University (Beijing, China). The expression plasmid for human ACE2 was obtained from GeneCopoeia (Guangzhou, China).

### Cell lines

HEK 293T cells were purchased from the American Type Culture Collection (ATCC, Manassas, VA, USA). Cells were maintained in Dulbecco’s modified Eagle medium (DMEM; Hyclone, Waltham, MA, USA) supplemented with 10% fetal bovine serum (FBS; Gibco, Rockville, MD, USA), 100 mg/mL of streptomycin, and 100 units/mL of penicillin at 37 °C in 5% CO2. HEK 293T cells transfected with human ACE2 (293T-ACE2) were cultured under the same conditions with the addition of G418 (0.5 mg/mL) to the medium. Elastase (from porcine pancreas) was obtained from MACKLIN Biochemical Co. (Shanghai, China).

### Antibodies and inhibitors

The anti-RBD monoclonal antibody against the SARS-CoV-2 S protein was kindly provided by Prof. Aishun Jin from Chongqing Medical University. Recombinant human ACE2 linked to the Fc domain of human IgG1 (ACE2-Ig) was purchased from Sino Biological Inc. Sivelestat sodium (MedChemExpress, Monmouth Junction, NJ, USA) Camostat mesylate (Tokyo Chemical Industry, Tokyo, Japan), and aloxistatin (E-64d; MedChemExpress) were dissolved in dimethyl sulfoxide (DMSO) at a stock concentration of 50 mM.

### Sera samples

A total of 70 convalescent sera samples from patients with COVID-19 (at 2–4 weeks after symptom onset) were collected from three designated hospitals in Chongqing from February 1 to February 10, 2020 (Supplementary information, Table S1). All sera were tested positive using magnetic chemiluminescence enzyme immunoassay (MCLIA) kits supplied by BioScience Co. (Tianjin, China)^29^. Patient sera were incubated at 56 °C for 30 min to inactivate the complement prior to experiments.

### SARS-CoV-2 genome analysis

The online Nextstrain analysis tool (https://nextstrain.org/ncov) was used to track the D614G mutation in SARS-CoV-2 genomes. All the 2,834 genomes sampled between Dec 20, 2019 and Jun 12, 2020 were visualized using the ‘rectangular’ layout. The mutations were labeled on branches.

All complete SARS-CoV-2 S gene sequences were downloaded from The National Center for Biotechnology Information (NCBI) website (https://www.ncbi.nlm.nih.gov/sars-cov-2/) on Jun 1, 2020. We obtained 4,701 S coding sequences from NCBI, and after excluding partial and frameshift sequences, 4,649 completed S sequences were used for further analysis. All S nucleotide sequences were translated to amino acid sequences. The nucleotide and amino acid sequences of S were aligned with multiple sequence alignment software MUltiple Sequence Comparison by Log-Expectation (MUSCLE) separately. The ‘Wuhan-Hu-1’ strain (NC_045512) was used to as the reference sequence, and the mutations were extracted using private PERL scripts.

### Western blot analysis of SARS-CoV-2 S protein expression

To analyze S protein expression in cells, S-D614- and S-G614-expressing plasmids were transfected into HEK 293T cells. Total protein was extracted from cells using radio immunoprecipitation assay Lysis Buffer (CoWin Biosciences, Beijing, China) containing 1 mM phenylmethylsulfonyl fluoride (Beyotime, Shanghai, China). Equal amounts of protein samples were electrophoretically separated by 10% sodium dodecyl sulfate polyacrylamide gel electrophoresis, and then transferred to polyvinylidene difluoride membrane (Millipore, Billerica, MA, USA). The immunoblots were probed with the indicated antibodies. Protein bands were visualized using SuperSignal West Pico Chemiluminescent Substrate kits (Bio-Rad, Hercules, CA, USA) and quantified by densitometry using ImageJ software (NCBI, Bethesda, MD, USA).

### Production and titration of SARS-CoV-2 S pseudoviruses

SARS-CoV-2 pseudotyped viruses were produced as previously described with some modifications^30^. Briefly, 5 × 10^6^ HEK 293T cells were co-transfected with 6 μg each of pNL4-3.Luc.R-E- and recombinant SARS-CoV-2 S plasmids (pS-D614 or pS-G614) using Lipofectamine 3000 Transfection Reagent (Invitrogen) according to the manufacturer’s instructions. The S-D614 and S-G614 protein pseudotyped viruses in supernatants were harvested 48 h after transfection, centrifuged, filtered through a 0.45 μm filter, and stored at −80°C. The pMD2.G was co-transfected with the pNL4-3.Luc.R-E-plasmid to package the VSV-G pseudovirus.

The titers of the pseudoviruses were calculated by determining the number of viral RNA genomes per mL of viral stock solution using RT-qPCR with primers and a probe that target LTR^31^. Sense primer: 5’-TGTGTGCCCGTCTGTTGTGT-3’, anti-sense primer: 5’-GAGTCCTGCGTCGAGAGAGC-3’, probe: 5’-FAM-CAGTGGCGCCCGAACAGGGA-BHQ1-3’. Briefly, viral RNAs were extracted using TRIzol (Invitrogen, Rockville, MD) and treated with RNase-free DNase (Promega, Madison, WI, USA) and re-purified using mini columns. Then, the RNA was amplified using the TaqMan One-Step RT-PCR Master Mix Reagents (Applied Biosystems, Thermo Fisher). A known quantity of pNL4-3.Luc.R-E-vector was used to generate standard curves. The S-D614 and S-G614 protein pseudotyped viruses were adjusted to the same titer (copies/mL) for the following experiments.

### SARS-CoV-2 S-mediated pseudoviral entry assay

To detect S variant-mediated viral entry, 293T-ACE2 cells (2 × 10^4^) grown on 96-well plates were infected with the same amount of S-D614 or S-G614 pseudovirus (3.8 × 10^4^ copies in 50 μL). The cells were transferred to fresh DMEM medium 8 h post-infection, and RLU was measured 24-72 h post-infection using Luciferase Assay Reagent (Promega, Madison, WI, USA) according to the manufacturer’s protocol^32^.

### Neutralization and inhibition assays

The 293T-ACE2 cells (2 × 10^4^ cells/well) were seeded on 96-well plates. For the neutralization assay, 50 μL pseudoviruses, equivalent to 3.8 × 10^4^ vector genomes, were incubated with serial dilutions of sera samples from patients and normal human serum as a negative control for 1 h at 37 °C, then added to the 293T-ACE2 cells (with three replicates for each dilution). For the inhibition assay, the cells were pretreated with elastase for 5 min and then infected with pseudotyped viruses in the presence of various concentrations of sivelestat sodium. After incubation for 12 h, the medium was replaced with fresh cell culture medium. Luciferase activity was measured 72 h after infection and the percentage of neutralization was calculated using GraphPad Prism 6.0 software (GraphPad Software, San Diego, CA, USA). Percentage of RLU reduction (inhibition rate) was calculated as: 1- (RLU of sample sera-control wells) / (RLU from mock control sera-control wells)) ×100%. The titers of neutralizing antibodies were calculated as 50% inhibitory dose (ID_50_).

### Statistical analyses

Statistical analyses of the data were performed using GraphPad Prism version 6.0 software. Quantitative data in histograms are shown as means ± SD. Statistical significance was determined using ANOVA for multiple comparisons. Student’s *t*-tests were applied to compare the two groups.

Differences with *P* values < 0.05 were deemed statistically significant.

### Ethical approval

The study was approved by the Ethics Commission of Chongqing Medical University (ref. no. 2020003). Written informed consent was waived by the Ethics Commission of the designated hospital for emerging infectious diseases.

## Acknowledgments

We would like to thank Prof. Cheguo Cai (Wuhan University, Wuhan, China) for providing the pNL4-3.Luc.R-E-plasmid. We also grateful to Dr. Hongbing Jiang (Washington University in St. Louis, St. Louis, MO, USA) for the suggestions. We gratefully acknowledge the authors, originating and submitting laboratories of the SARS-CoV-2 sequence data from GISAID. This work was supported by the Emergency Project from the Science & Technology Commission of Chongqing (cstc2020jscx-fyzx0053 to A-L.H.), the Emergency Project for Novel Coronavirus Pneumonia from the Chongqing Medical University (CQMUNCP0302 to K.W.), the Leading Talent Program of CQ CSTC (CSTCCXLJRC201719 to N.T.), and a Major National Science & Technology Program grant (2017ZX10202203 to A-L.H.) from the Science & Technology Commission of China.

## Author contributions

A-L.H., N.T., and K.W. conceived the project and supervised the study. J.H., C-L.H., Q-Z.G. and G-J.Z. performed most experiments. J.H. and K.W. performed serum neutralization assay. H-J.D. performed SARS-CoV-2 genome analysis. L-Y.H. performed structural analysis of S protein. Q-X.L., J.C. and X-X. C. collected the serum samples. J.H. and Q-Z.G. contributed to the statistical analysis. K.W. and N.T. wrote the manuscript. All the authors analyzed the final data, reviewed and approved the final version.

