## Supplemental Figure 1 for "D614G mutation of SARS-CoV-2 spike protein enhances viral infectivity"

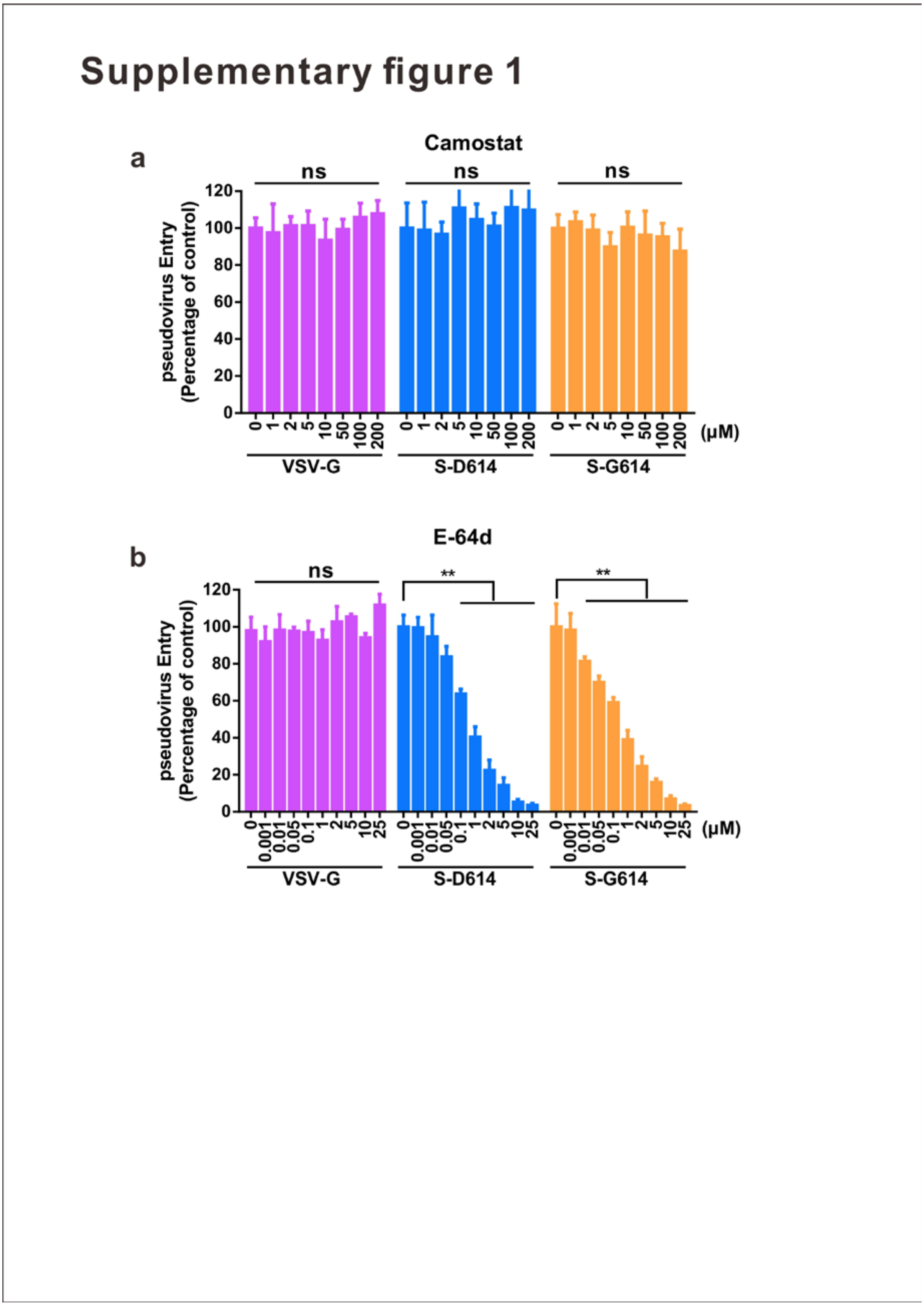


**Supplementary information, Fig. S1 Detection of protease inhibitors against SARS-CoV-2 pseudoviral infection. a-b,** Transmembrane protease serine 2

(TMPRSS2), cathepsin (Cat) B/L, and elastase-2 inhibitors were evaluated by pseudoviral cell entry assay. 293T-ACE2 (human angiotensin-converting enzyme 2) cells were pre-incubated with camostat mesylate (**a**) or E-64d (**b**), and subsequently inoculated with pseudoviruses. The VSV-G pseudovirus was used as the control. RLU was measured at 72 h post-pseudovirus inoculation. Cell viability was examined using MTT (3-(4,5-dimethylthiazol-2-yl)-2,5-diphenyltetrazolium bromide) assay. n=3, ±SD. **$P < 0.01$. ns, not significant.
